# TCIApathfinder: an R client for The Cancer Imaging Archive REST API

**DOI:** 10.1101/240986

**Authors:** Pamela Russell, Kelly Fountain, Dulcy Wolverton, Debashis Ghosh

## Abstract

**Summary:** The Cancer Imaging Archive (TCIA) hosts publicly available de-identified medical images of cancer from over 25 body sites and over 30, 000 patients. Over 400 published studies have utilized freely available TCIA images. Image series and metadata are available for download through a web interface or a REST API. We present TCIApathfinder, an R client for the TCIA REST API. TCIApathfinder wraps API access in user-friendly R functions that can be called within an R session or easily incorporated into scripts. Functions are provided to explore the contents of the large database and to download image files.

**Availability and implementation:** TCIApathfinder is available under the MIT license as a package on CRAN (https://cran.r-project.org/web/packages/TCIApathfinder/index.html) and at https://github.com/pamelarussell/TCIApathfinder.

## Introduction

### Radiomics

The process known as radiomics involves the conversion of digital medical images into higher-dimensional data and the subsequent mining of these data (Kumar *et al*., 2012). Hundreds or even thousands of image features can be extracted from radiological image analysis and associated with biologic or clinical endpoints to develop diagnostic, prognostic and predictive models (Avanzo *et al*., 2017). While this can be applied to many biomedical areas, oncologic applications are of particular interest.

Oncologic treatment failure is often attributed to heterogeneity within tumors at the phenotypic, physiologic, and genomic levels (Gerlinger *et al*., 2012; Sequist *et al*., 2011; Sottoriva *et al*., 2013; Yachida *et al*., 2010). A major aim of radiomics is to provide quantitative measurements of intra- and inter-tumoral heterogeneity, thereby individualizing treatment (Gillies *et al*., 2016). Unlike tissue biopsy which offers a small sample of the tumor for analysis, radiomics has the potential to evaluate the whole tumor in the native environment. Unfortunately, there are multiple challenges to the process of radiomics. The need for data and ability to share data has been cited as one of the largest hurdles to advancing the field of radiomics (Avanzo *et al*., 2017; Gillies *et al*., 2016). Large centralized data repositories such as The Cancer Imaging Archive (TCIA) offer a solution.

### Overview of TCIA

TCIA provides de-identified clinical images of cancer for use by the research community (Clark *et al*., 2013). Currently, TCIA includes images from over 25 cancer types and over 30,000 patients. TCIA also hosts supporting data related to the images, as well as analysis results from the research community based on TCIA data. TCIA is contributing to research efforts toward understanding the genomic basis of cancer by providing over 20 collections of clinical images from patients whose matched tumor genomic profiles are freely available from The Cancer Genome Atlas (Cancer Genome Atlas Research Network *et al*., 2013).

### TCIA data

TCIA datasets, referred to as “collections”, typically represent sets of patients sharing a common disease. Descriptions of each collection are available from the TCIA website. A variety of imaging modalities are represented, such as magnetic resonance imaging (MRI), computed tomography (CT), and positron emission tomography (PET). The image files in TCIA conform to the widely adopted DICOM standard (Mustra *et al*., 2008). For each patient, one or more image studies are included. A study may include one or more image series. In turn, an image series is a stack of two-dimensional images from a single run of an instrument. Patient names and birth dates have been de-identified; patient sex and age are provided. TCIA supports reproducibility through the use of Digital Object Identifiers (DOIs) to refer to subsets of data.

## Description of TCIApathfinder

TCIApathfinder can be installed directly from the Comprehensive R Archive Network (CRAN) or from https://github.com/pamelarussell/TCIApathfinder. Package documentation is available as a PDF manual from CRAN, from within an R session using R’s documentation system, or on GitHub. In order for the package to function correctly, an API key must be obtained from TCIA.

In TCIApathfinder, function calls are used to explore the available data in TCIA and to download image files to the local machine. Two functions, *save_single_image()* and *save_image_series*(), download image files from TCIA to the local machine. The remaining functions in the package are used to explore the available data in TCIA. Each exploratory function returns an object containing simplified summarized data, a parsed JSON response, and the raw API response. Functions to obtain available values for metadata features include *get_collection_names()*, *get_modality_names()*, *get_body_part_names()*, and *get_manufacturer names()*. Each of these can be optionally restricted to subsets of the data by other metadata features. Patient information is retrieved with *get_patient_info()* and *get_patients_by_modality()*, both of which can be restricted to a particular collection. Individual image study IDs are obtained with *get_studies_in_collection()*, which can be restricted to a particular patient. More extensive metadata about image studies — at the collection, patient, or study level — is returned by *get_patient_studies()*. Information on image series is returned by *get_series_info()*, which can be restricted by many possible metadata values, and *get_series_size()*. Individual DICOM image IDs within an image series are obtained with *get_sop_instance_uids()*. Recent additions to TCIA can be listed with *get_new_patients_in_collection()* and *get_new_studies_in_collection()*.

TCIApathfinder can be loaded into an active R session and used directly from the R console, or incorporated into R scripts.

## Conclusion

TCIApathfinder makes the extensive resources in TCIA easily available and accessible in the highly popular R programming environment. Simple functions allow the full collection to be easily explored before patients are selected for analysis. Images and supporting data can be imported directly into R scripts for further analysis using packages such as oro.dicom (Whitcher *et al*., 2011) and other image analysis packages. For patients also included in The Cancer Genome Atlas, tumor genomic data can be imported into R via the matched patient ID using the TCGAbiolinks package (Colaprico *et al*., 2016) and analyzed using the extensive tools in Bioconductor (Gentleman *et al*., 2004). Vignettes included with the package demonstrate TCIApathfinder usage as well as downstream analysis with other packages. TCIApathfinder will significantly lower the barrier for researchers to leverage the valuable resources in TCIA.

## Acknowledgements

We thank Bernard Jones and Julio Carballido-Gamio for valuable conversations about medical image analysis and the DICOM standard.

## Funding

This work has been supported by the Grohne-Stapp Endowed Chair for Cancer Research (University of Colorado Cancer Center).

## References

Avanzo, M. et al. (2017) Beyond imaging: The promise of radiomics. Phys. Med., 38, 122–139.

Cancer Genome Atlas Research Network et al. (2013) The Cancer Genome Atlas Pan-Cancer analysis project. Nat. Genet., 45, 1113–1120.

Clark, K. et al. (2013) The Cancer Imaging Archive (TCIA): maintaining and operating a public information repository. J. Digit. Imaging, 26, 1045–1057.

Colaprico, A. et al. (2016) TCGAbiolinks: an R/Bioconductor package for integrative analysis of TCGA data. Nucleic Acids Res., 44, e71.

Gentleman, R.C. et al. (2004) Bioconductor: open software development for computational biology and bioinformatics. Genome Biol., 5, R80.

Gerlinger, M. et al. (2012) Intratumor heterogeneity and branched evolution revealed by multiregion sequencing. N. Engl. J. Med., 366, 883–892.

Gillies, RJ. et al. (2016) Radiomics: Images Are More than Pictures, They Are Data. Radiology, 278, 563–577.

Kumar, V. et al. (2012) Radiomics: the process and the challenges. Magn. Reson. Imaging, 30, 1234–1248.

Mustra, M. et al. (2008) Overview of the DICOM standard. In, 2008 50th International Symposium ELMAR., pp. 39–44.

Sequist, L.V. et al. (2011) Genotypic and histological evolution of lung cancers acquiring resistance to EGFR inhibitors. Sci. Transl. Med., 3, 75ra26.

Sottoriva, A. et al. (2013) Intratumor heterogeneity in human glioblastoma reflects cancer evolutionary dynamics. Proc. Natl. Acad. Sci. U. S. A., 110, 4009–4014.

Whitcher, B. et al. (2011) Working with the DICOM and NIfTI Data Standards in R. Journal of Statistical Software, Articles, 44, 1–29.

Yachida, S. et al. (2010) Distant metastasis occurs late during the genetic evolution of pancreatic cancer. Nature, 467, 1114–1117.

